# Structural Insights into Cir-mediated Killing by the Antimicrobial Protein Microcin V

**DOI:** 10.1101/2025.03.06.641670

**Authors:** Stavros A. Maurakis, Angela C. O’Donnell, Istvan Botos, Rodolfo Ghirlando, Bryan W. Davies, Susan K. Buchanan

## Abstract

Drug-resistant bacterial infections have become a high-priority concern in the global healthcare landscape. Novel treatments for such infections are needed, but can be especially difficult to develop for Gram-negative species owing to the need to traverse the outer membrane to reach targets beneath. A promising solution is found in natural antibiotics such as albomycin, which can bind outer membrane siderophore receptors and co-opt them for import into the periplasm. Exploring this and similar mechanisms may open avenues for antibiotic development. An underappreciated class of natural antibiotics are the microcins, which are small antimicrobial proteins secreted by certain bacterial species as a means of inter-species competition. Once secreted, the microcins bind specific outer-membrane receptors of prey species and cross into the periplasm. Microcins have potent activity, bind specific targets, and have been shown to control pathobiont expansion and pathogen colonization in animal models. One such microcin, MccV, has previously been shown to utilize the *E. coli* colicin Ia receptor, Cir, for periplasmic import. Here, we report the first high-resolution structure of the Cir/MccV complex by Cryo-EM, revealing an interaction centered on an electropositive cavity within a usually occluded binding pocket in Cir. We also report the calculated affinity of MccV for Cir. We mutagenized putative interacting residues at this interface and identified key contacts that are critical to MccV binding and Cir-mediated import and bacteriolysis. Future efforts in this project will help us better understand the mechanisms of microcin killing, and will assess relationships between other microcins and their respective importers with the aim of understanding the potential for microcins to be used as treatment for drug-resistant pathogens.

## INTRODUCTION

Increasing incidence of drug-resistant bacterial infections represents a growing threat to global health^1,2^. Notably among these, Gram-negative species make up four of the six ESKAPE pathogens^3^, and while not listed as an original member of these six, the ubiquitous Gram-negative *Escherichia coli* has also demonstrated a marked increase in antimicrobial drug resistance (AMR) among certain isolates^4^. To be effective, drugs must either traverse the Gram-negative outer membrane to access targets inside or target the membrane directly; both approaches pose obstacles to drug development or clinical deployment^5–8^. Overcoming the barrier to drug uptake would represent an important step forward in drug development. One approach is to take advantage of the innate outer-membrane transport systems already present in the bacteria, such as the Ton system^9,10^. Indeed, antibiotics which bind bacterial siderophores, common substrates of TonB-dependent transporters (TBDTs), and co-opt them for import have been described^11–14^. While most of the efforts in this arena have been limited to siderophore conjugates, an understudied class of bacteriocins called microcins represent another potential target.

Microcins are small (<10 kDa) antibacterial proteins which, when secreted by producer bacteria, localize to and bind specific receptors on the outer-membrane of other bacterial targets, including TBDTs^15–17^ (Figure 1A-B). Once imported, microcins block key functions in the target cell such as inhibiting protein synthesis, inducing double-stranded DNA breakage, or impairing ATP synthesis. Others kill more directly by forming pores in the bacterial membrane^15,18,19^. Harm to producer strains is prevented by production of an immunity protein^20^. Broadly, the microcins are divided into class I and class II; class I members are highly post-translationally modified while class II are largely unmodified except for the permissible presence of disulfide bonds (class IIa) or C-terminal attachment of iron siderophores (IIb)^21–23^. Besides their promising import phenotypes, the microcins demonstrate several other characteristics which suggest they may have strong applications in combatting AMR: they are highly potent, are tolerant of a large pH range, and do not show toxicity to human cells^24–26^.

**Figure 1.**
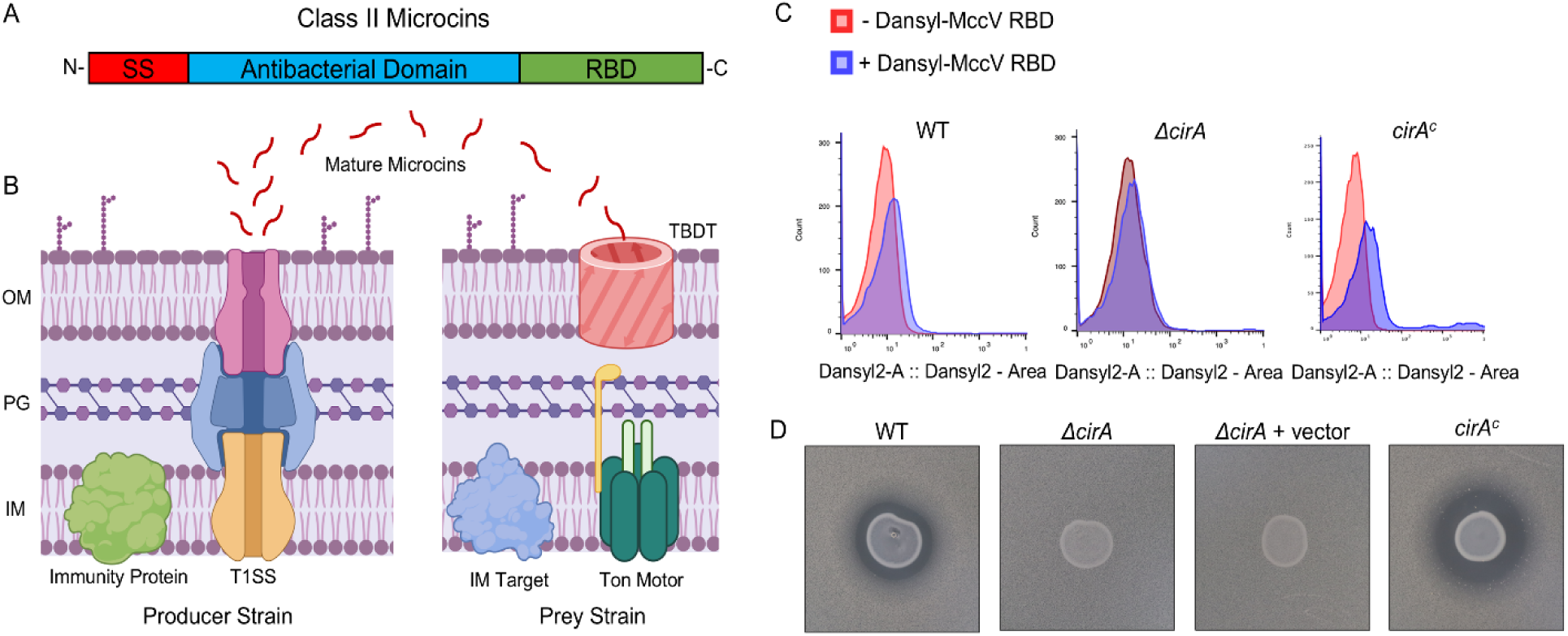
Class II Microcins Interact with TonB-dependent Transporters. A) Schematic representation of Class II microcin domains. SS: Signal Sequence, RBD: Receptor Binding Domain. B) Schematic overview of class II microcin function. Immature pre-microcins are produced in the cytoplasm of producer strains and subsequently exported by Type I secretion (T1SS). Mature microcins in supernatant then localize to the surface of prey bacteria and are imported by exploiting receptors from the Ton (and Tol) systems. Once they have entered the periplasm, microcins are free to interact with their cognate targets at the inner membrane. Producer strains are protected from auto-bactericidal activity via production of an immunity protein. OM: Outer Membrane, PG: Peptidoglycan, IM: Inner Membrane, TDBT: TonB-dependent Transporter. C) Flow cytometry histograms showing dansyl fluorescence shifts. WT *E. coli* cells treated with dansyl-MccV RBD (blue peaks) exhibit a shift relative to the untreated population (red peaks), and a greater shift is observed when Cir is overexpressed from a plasmid (right graph). This shift is lost when the Cir receptors are absent in a *cirA*::kan mutant (middle graph). D) Zone of inhibition assays demonstrating the sensitivity of *E. coli* to spotted predator strain secreting WT MccV. The WT *E. coli* lawn demonstrates clearance indicating sensitivity to MccV. This activity is lost when *cirA* is knocked out. Activity is restored when a plasmid expressing Cir is transformed into the *cirA*::kan strain.

The class IIa MccV (Microcin V, formerly Colicin V), produced mainly by *E. coli* but found in certain other *Enterobacteriaceae*, is well characterized among the microcins^27–29^. Once exported, MccV exhibits bactericidal activity against various *Escherichia, Salmonella, Shigella,* and *Klebsiella* species^30^. MccV is recognized by Cir, an iron-regulated TonB-dependent siderophore receptor that is also the target of the antimicrobial Colicin Ia^31–33^. Upon Cir-dependent import to the periplasm, MccV targets the inner-membrane serine transporter SdaC, ultimately resulting in collapse of membrane potential due to membrane pore formation^34,35^. However, neither the specific nature of MccV recognition of Cir, nor its interactions with the Ton motor (TonB, ExbB, ExbD), have previously been described.

To understand how MccV binds Cir, we solved the first structure of this complex via cryogenic electron microscopy (cryo-EM). Combined with binding assays utilizing purified biomolecules, we report that Cir and MccV bind with affinity in the low micromolar range, and that the structural nature of the binding interaction is conserved between known Cir ligands. Lastly, using site-directed mutagenesis, we interrogated the Cir/MccV binding and import mechanism and identified critical amino acid residues responsible for binding and effective import and bactericidal activity by MccV. Ultimately, these findings provide a critical advancement in the understanding of microcin biology and represent an important step toward characterizing the microcins as potential treatments against high-priority AMR bacteria.

## RESULTS

### MccV Binds and Kills Cir-expressing *E. coli*

Previous reports indicate that MccV is recognized by Cir, and the interaction results in bacterial death from disruption of membrane potential and an as-yet-undescribed interaction with the inner-membrane serine transporter SdaC^31,34,35^. However, little is known about the exact nature of their binding interaction or the structural nature of MccV docking with Cir. As such, we sought to interrogate these topics to gain further insights into microcin form and function.

Previous research demonstrated that the 32 C-terminal amino acids of MccV were sufficient to confer specificity to Cir for microcin cell entry^36^. To validate this genetic finding, we first established whether the C-terminal receptor binding domain (RBD) of MccV alone was capable of binding Cir-expressing *E. coli*. To this end, we synthesized a dansyl-tagged peptide comprised of the 32 RBD residues of MccV and tested for binding using flow cytometry (Figure 1C). This peptide caused a fluorescence shift in a wildtype culture of *E. coli* but not in the isogenic *cirA* deletion strain. This effect was complemented by expression of *cirA* from a plasmid. From here, we set out to define a bactericidal assay which demonstrated Cir-dependent bacteriolysis by secreted MccV. We demonstrated that when *E. coli* producing both MccV and its immunity protein are spotted on a lawn of sensitive *E. coli*, a clear zone of inhibition is observed, indicating MccV antibacterial action against the sensitive strain. This zone was lost when *cirA* was deleted, while sensitivity was restored by *cirA* expression *in trans*. (Figure 1D). This killing assay gave us a framework within which to probe phenotypic changes as we further characterized the interaction between the proteins.

### Structure Determination of the Cir/MccV Complex

4-Met substituted *E. coli* Cir^32^ was expressed into native membranes of *E. coli* strain BL21 (DE3) which were then isolated. The membrane fraction was incubated with purified MccV and then solubilized using n-dodecyl-β-D-maltoside (DDM) and purified using a 10X N-terminal histidine tag on Cir. After affinity chromatography, DDM was replaced with nonionic amphipol (NAPol)^37^ as a stabilizing agent for Cir, and the remaining purification steps were performed in the absence of detergent. The NAPol-stabilized Cir/MccV complex was then run over size exclusion chromatography. Under these conditions, Cir and MccV co-eluted cleanly as a single peak. Cryo-EM analysis of these fractions showed homogenous, well-distributed particles which were used for data collection and single particle analysis. (Supplementary Figure 1).

In total, 102,419 particles were used to calculate the three-dimensional (3D) map of the Cir/MccV complex in NAPol at 2.9 Å resolution (Figure 2, Supplementary Figure 2, Table 1). A predicted model was generated using AlphaFold2^38^ and fit into the density map, then manually adjusted for best fit. The entire structure was then refined to a final map-to-model correlation coefficient of 0.88. Our cryo-EM structure of Cir shows the prototypical architecture of a TBDT, including a 22-stranded β-barrel occluded by an N-terminal plug, and 11 extracellular loops of variable length^39^. More specifically, it has overall dimensions consistent with those previously observed during crystallographic analysis of the Cir apo-form (PDB code 2HDF): an ellipsoid β-barrel approximately 40 Å high with inner dimensions of approximately 40 Å across the short axis and 50 Å across the long axis, and extracellular loops 7 and 8 being the longest, occupying most of the area across the top of the barrel^32^ (Figure 2A-B). Loop 2, made up of residues 210-231, was not resolved in the original crystal structure but was visible in the cryo-EM map. This loop is the third largest and sits on the opposite side of the β-barrel from loops 7 and 8.

**Figure 2.**
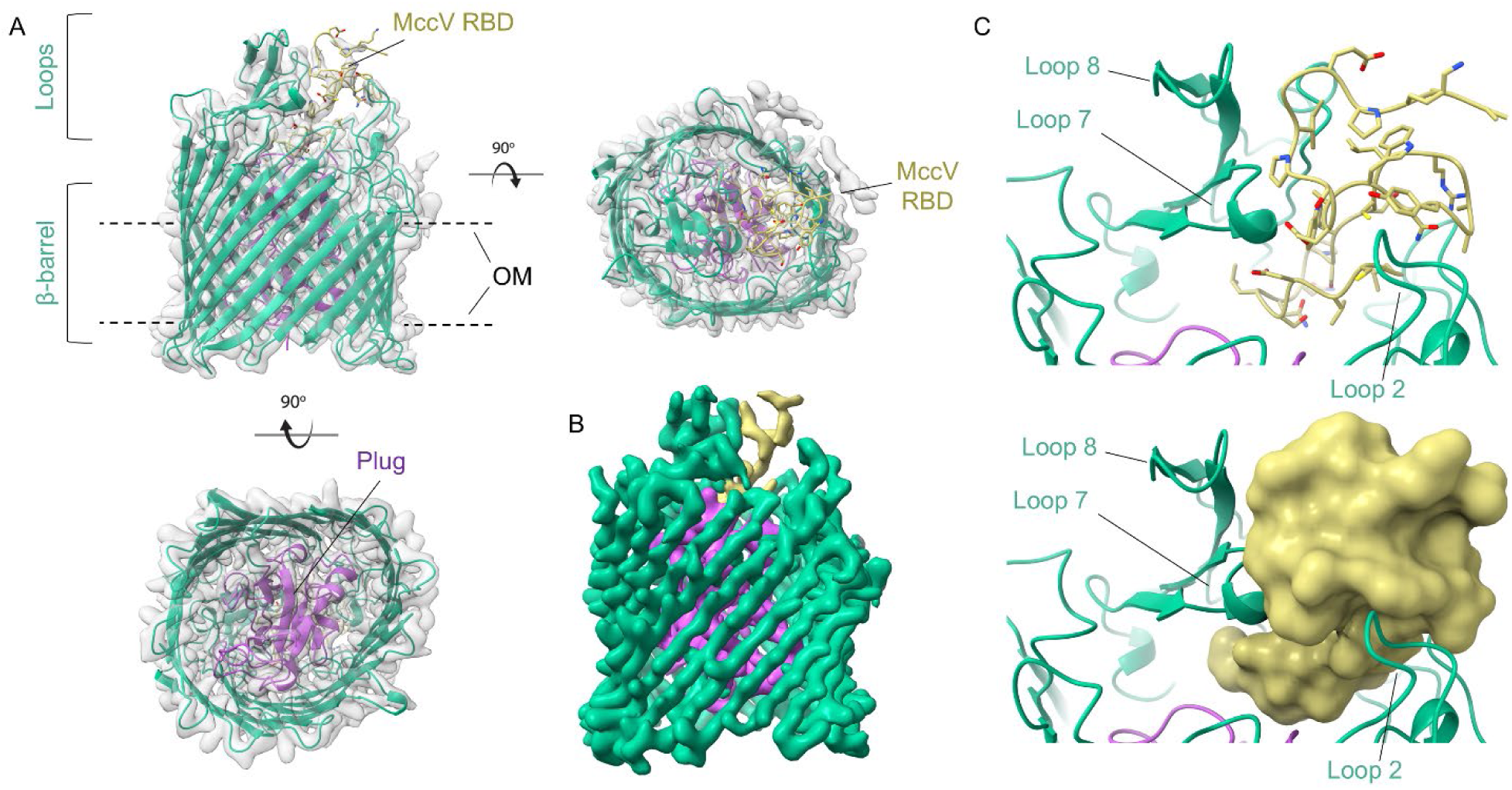
Structure of Cir Bound by the MccV Receptor Binding Domain. A) Cryo-EM structure of the Cir/MccV RBD complex. The β-barrel and extracellular loops of Cir are shown as green ribbon with the plug domain highlighted in purple. The MccV RBD is shown in gold. The structure is inset into the cryo-EM density map with the NAPol micelle subtracted. OM = outer membrane. B) Cir/MccV RBD cryo-EM map colored to match model (contour level 0.120). C) Illustration of the Cir/MccV binding pocket shown as cartoon (top) and as MccV surface (bottom). MccV interacts with the Cir extracellular loops in an area flanked by loops 2, 7, and 8 at the top and the plug domain below. This map and accompanying coordinates have been deposited to EMDB and PDB under codes EMD-49565 and 9NN6, respectively.

While mature MccV consists of residues alanine 16 to leucine 103 (residues 1-88 by our numbering), the cryo-EM map showed strong signal only for residues 57-88, which correspond to the C-terminal RBD (Figure 2B). Inspection of this section of the map showed that the MccV RBD settles into a binding pocket atop the plug domain and flanked on one side by Cir loops 7 and 8, and on the other by loop 2 (Figure 2C). In total, an interacting surface of 1232.8 Å^2^ is buried during complex formation, representing 44.7% of the total MccV surface found in the model. Strikingly, this is the same space occupied by the Colicin Ia R-domain upon its docking with Cir^32^.

### MccV Binds Cir Similarly to Colicin Ia

Upon observing that MccV bound Cir in the same position as Colicin Ia, we wondered whether there were similarities in their binding modes and mechanisms as well. When compared to apo-Cir, we observed a large deflection away from the top of the plug by loops 7 and 8, with the tip of each loop moving 11.5 Å and 12 Å, respectively, from its position in the apo-form (Figure 3A, Movie 1). This change in conformation reveals the binding pocket and opens a space for MccV to occupy, which would otherwise be occluded (Figure 3B). Parts of Cir loop 2 also show close proximity to MccV, with sidechains from each protein separated by as little as 3.5 Å. Comparison of the apo- and ligand-bound structures shows an outward twisting movement at the base of this loop, suggesting that it too may be displaced during binding, but because this loop was not well resolved in the apo-form crystal structure, we could not assess this fully. We also observed slight movement in the residues corresponding to the Cir TonB Box^40^ (T32-A37, Movie 2), suggesting a possible adaptation of a plug conformation which allows access and binding by TonB. This is consistent with previous reports which showed the necessity of TonB for MccV-dependent bactericidal activity^31^.

**Figure 3.**
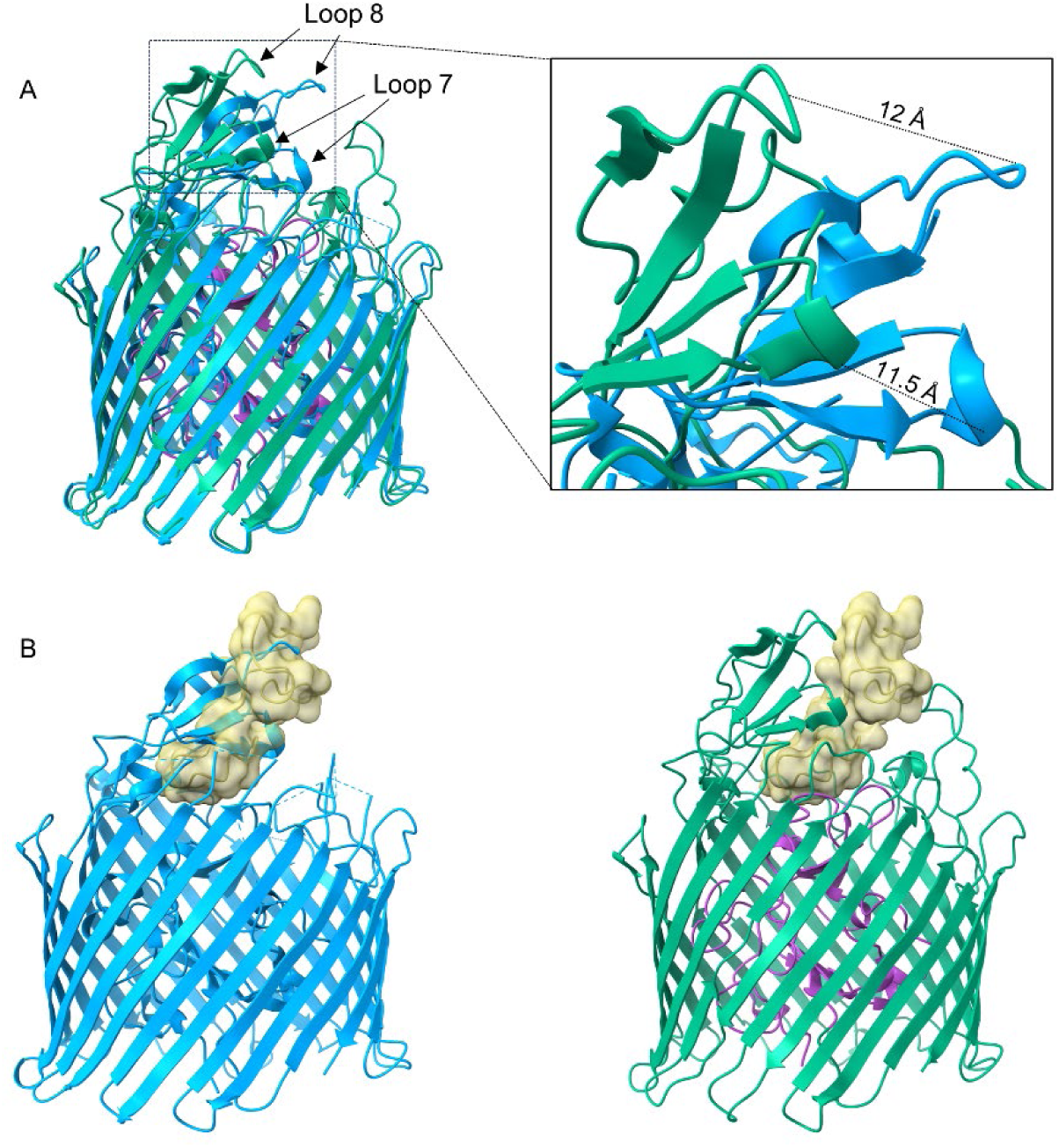
Cir Undergoes Conformational Changes to Facilitate MccV Docking. A) Cir/MccV RBD (MccV RBD not shown) superimposed onto apo-Cir (PDB code 2HDF, blue). Loops 7 and 8 from the complex structure are displaced relative to their positions in the ligand-free structure by 11.5 Å and 12 Å, respectively. B) MccV overlaps the space vacated by loops 7 and 8. Left: apo-Cir with MccV RBD (cartoon inset into surface, gold) superimposed. MccV occupies the same space as loops 7 and 8 from the apo structure. Right: The complex structure with loops 7 and 8 deflected to facilitate MccV docking.

The rearrangement described above mimics the action of loops 7 and 8 upon Colicin Ia binding, which results in an even greater displacement: 17 Å in loop 7 and 21 Å in loop 8. Because MccV occupies the same space and causes approximately the same conformational changes to Cir as does Colicin Ia, we hypothesized that the nature of the binding interactions may be similar. The H-bonding network between Colicin Ia and Cir is well understood. Of note, R436 of Cir was shown to interact with D350 of Colicin Ia, with Colicin E369 also in proximity. Likewise, Colicin residues D358 and E357 are in proximity to Cir R490 and R116, respectively^32^. We noted Cir R116, R436, and R490 in our own model and probed the interaction of these side chains with MccV. Strikingly, these residues appear to form an electropositive cavity at the base of the loop 7/loop8 area above the plug, with the sidechain of MccV D85 settled into the middle (Figure 4A-B, Supplementary Figure 3). When in complex, charged atoms from D85 are located 3 Å, 4.3 Å, and 2.8 Å from complementary charges on R116, R436, and R490, respectively (Figure 4C-D). Further, we note a 2.8 Å salt bridge formed between OD1 from D85 and NH2 from R490. To further probe the Cir/MccV interaction, we performed clustering analysis using PICKLUSTER^41^. This analysis reinforced our supposition that the arginine cavity coordinated MccV binding, suggesting an interaction network anchored by R436 and R490, with contributions from C435 and S464. An additional minor interface was predicted between Cir R513 and MccV E61. With putative interacting residues identified, we set out to validate these findings via functional and binding analyses.

**Figure 4.**
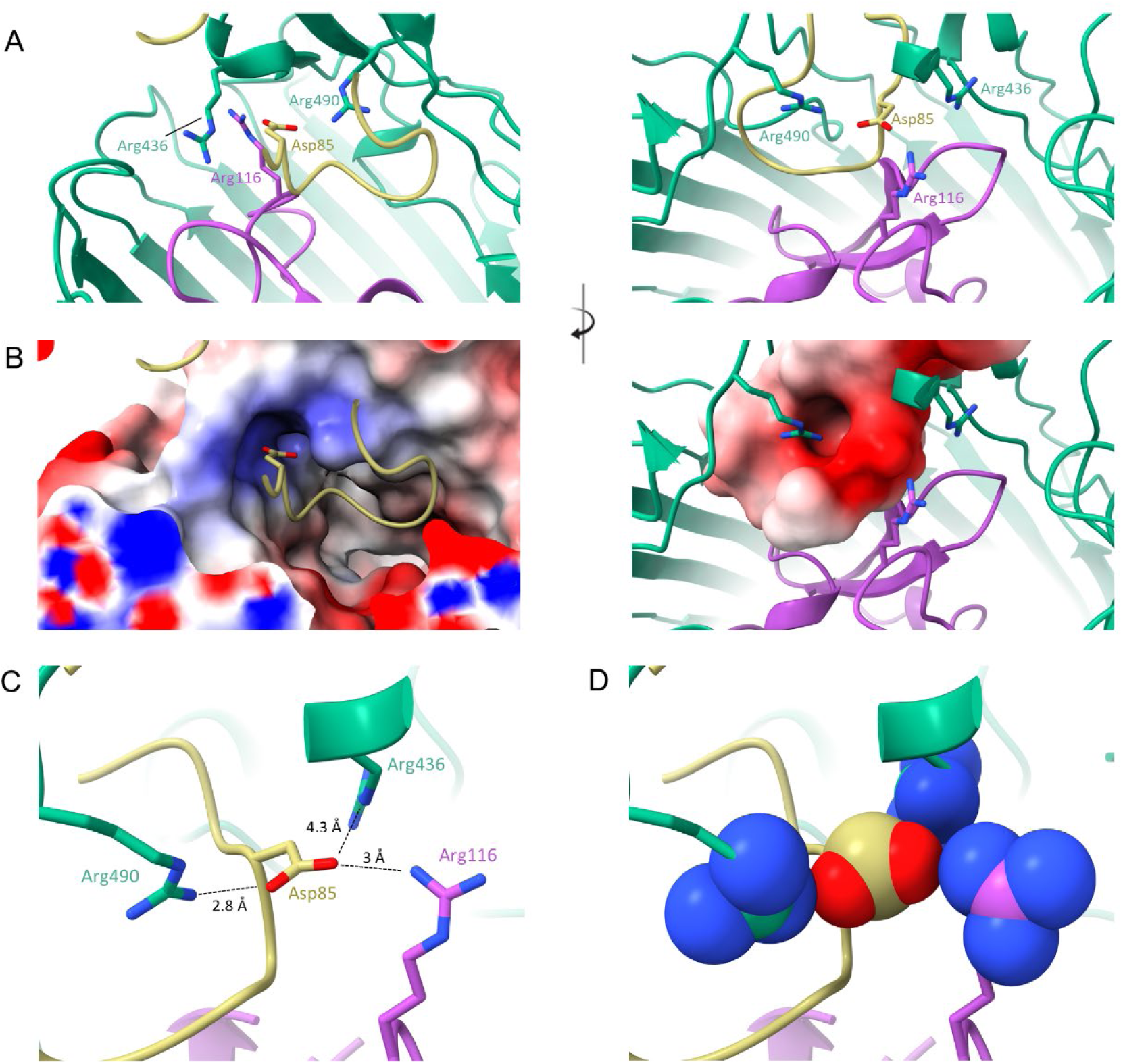
MccV Interacts with an Electropositive Pocket In the Cir Loops. Three arginine residues (Arg 436 from loop 7, Arg 490 from loop 8, and Arg 116 from the plug) form an electropositive cavity within the Cir ligand-binding pocket. When in complex, the sidechain of aspartate 85 from MccV fills the space inside the cavity, creating a putative electrostatic interaction. A-B) A shot-reverse shot illustration of the interacting residues shown as cartoon (top) and with electrostatic surface colored (bottom). Arginine cavity electropositive surface is shown on the left of panel B, with the electronegative Asp 85 surface shown on the right. C-D) Asp 85 settles in close proximity to the Cir aspartates. Atomic distances between the relevant atoms of Asp 85 and each of the arginines were determined and are illustrated in panel C, with sidechain heteroatom surfaces shown in D. Interactions shown: R116 NH1 with D85 OD2, R436 NE with D85 OD2, and R490 NH2 with D85 OD1.

### Mutagenesis Impairs MccV Binding, Import, and Bactericidal Function

We generated a series of mutations in Cir and MccV to probe their effects on MccV binding and function, with a primary focus on alanine substitution and swapping charged residues to the opposite charge. Cir mutants were stable and consistent with the wildtype when expressed in *E. coli* and mutations in both proteins did not affect purification (Supplementary Figure 4). We first assessed whether the mutations impacted Cir sensitivity to MccV killing. To this end, we performed zone of inhibition assays comparing MccV sensitivity between WT and *cirA* mutant constructs. We observed a range of loss of sensitivity in the C435A, R436A/E, R490A/E, and R436E/R490E Cir variants, as well as in D85A and D85R MccV (Figure 5A). We hypothesized that the impaired MccV function was concomitant with diminished or abolished binding, so we next set out to evaluate potential binding deficiencies introduced by our mutations. We first calculated the affinity between the wildtype proteins using biolayer interferometry (BLI). Using purified, biotinylated MccV, we loaded streptavidin coated BLI sensors which were then probed with dilutions of Cir, resulting in a dose-dependent response trace (Figure 5B). When fitted to a single site binding model, these data returned a calculated KD of 1.7 +/- 1.2 μM. Because this is the first time reporting the affinity of these partners, and because we planned to deploy this BLI-based approach for further characterization, we elected to validate our initial findings using another independent method. For this, we utilized sedimentation velocity analytical ultracentrifugation (SV-AUC). SV-AUC analysis of NAPol-stabilized Cir and its mixtures with MccV resulted in a single-site model KD of 0.6 +/- 0.2 μM, within the margin of error of our BLI results (Supplementary Figure 5). Based on concurrence of two experimental approaches, we confidently report a KD of approximately 1 μM with 1:1 binding stoichiometry. With a baseline binding affinity established, we purified both MccV mutants and the Cir variants with altered MccV sensitivity and performed the same BLI analysis for each (Figure 5C). D85A and D85R showed similar reductions in Cir binding relative to WT, with approximately a 40-45% reduction in response shift for the same probe concentration of 5 μM. There was more variation from the Cir mutations. C435A showed the smallest change and reached a peak response shift identical to that of the WT, albeit with a slower on-rate. R436E/R490E showed a moderate loss of approximately 55% response shift compared to WT. The most pronounced defect was observed in R490A, in which binding was almost completely abolished and response shift was similar to that observed when no probe was added.

**Figure 5.**
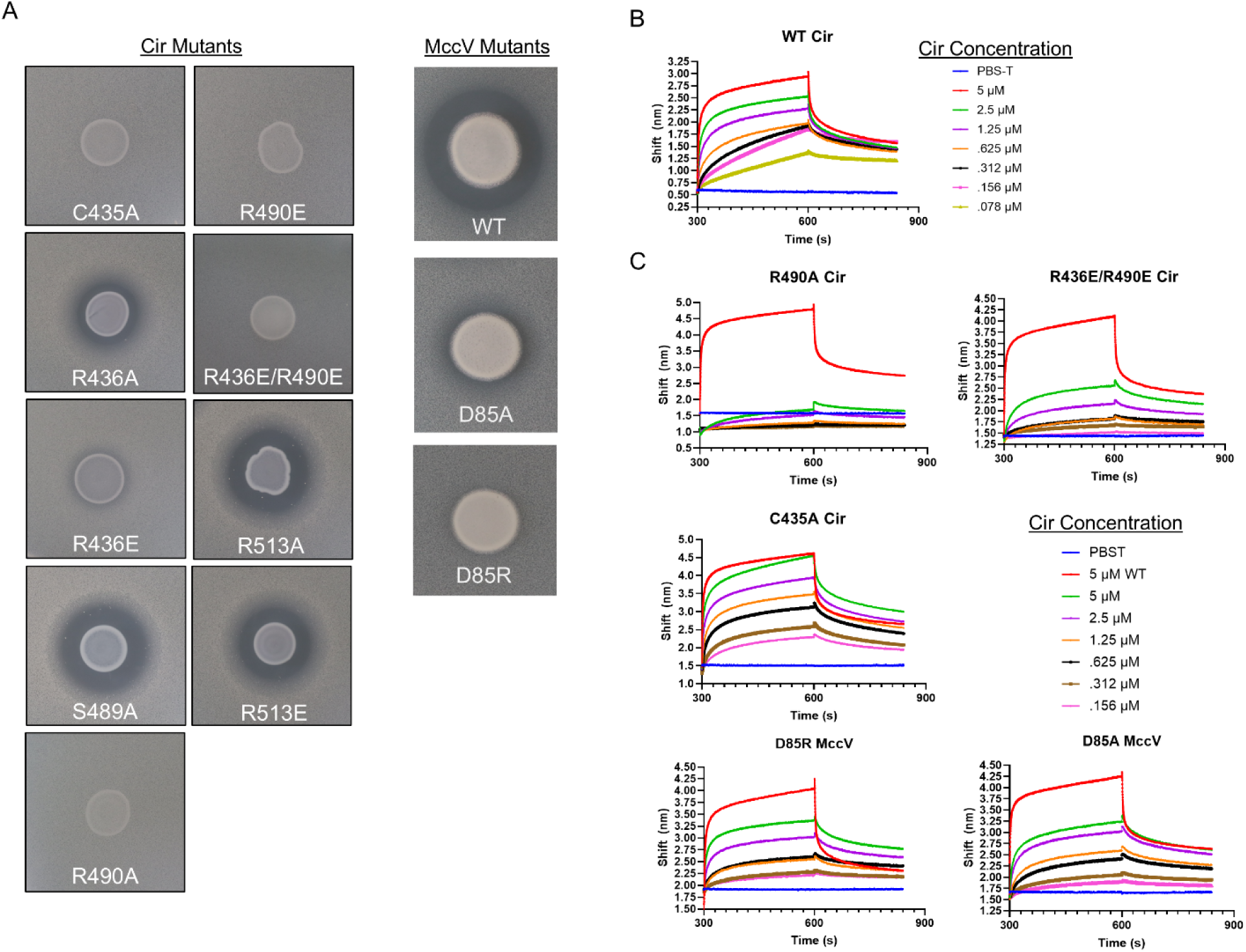
Binding Pocket Mutants Show Decreased Bactericidal Activity and Binding. A) Zone of inhibition assays demonstrating sensitivity of Cir mutants to WT MccV. Cir mutations C435A, R436E, R490E, R490A, and R436E/R490E exhibit losses in sensitivity to WT MccV secreted from *E. coli*. The third column shows that the toxic activity of MccV is attenuated when residue D85 is mutated. B) MccV binds Cir with high affinity. Purified MccV was biotinylated and loaded onto streptavidin-coated biolayer interferometry (BLI) sensors at a concentration of 100 nM, then probed with a dilution series (5 μM -> 78 nM in twofold steps) of wildtype (WT) Cir to determine affinity. C) Mutant binding assays. 100 nM WT or mutant biotinylated MccV was loaded onto BLI sensors and probed with dilutions of WT or mutant Cir to compare binding against the WT partners. In each mutant assay, 5 μM WT Cir was included as a positive control (red trace). For all BLI experiments, buffer alone (PBS-T) was added as a negative control. Representative BLI traces are shown from n=3 experiments.

Finally, we considered the role of the Ton motor on MccV import after initial binding to Cir at the cell surface. As mentioned, we observed a slight movement in the Cir residues comprising the TonB Box (Movie 2), and it has previously been shown that inactivation of the *tonB* and *exbB* genes prevented MccV uptake^31^. In addition, TonB Box point mutations which affect its crosslinking pattern with TonB are known to cause TonB uncoupling and subsequent loss of the import phenotye^42^. With this in mind, we utilized a Cir TonB Box mutant (M33C/V35P, Figure 6A) in both our agar plate killing assay and BLI to confirm that Cir, via the Ton motor, was the primary means of MccV traversal through the outer membrane (Figure 6B-C). When M33C/V35P-expressing cells were grown in the presence of MccV-secreting cells, we observed no zone of inhibition in the secretion area, indicative of no successful MccV import. Conversely, when we added M33C/V35P to BLI plates with WT MccV, we observed no reduction in binding signal compared to WT Cir at the same dilution, suggesting initial binding prior to import was unaffected. Collectively, these observations confirm the role of the electropositive cavity in the Cir binding pocket as a key target for MccV binding, and we conclude that Cir, via TonB-dependent import, is also the key translocation path for MccV into the periplasm.

**Figure 6.**
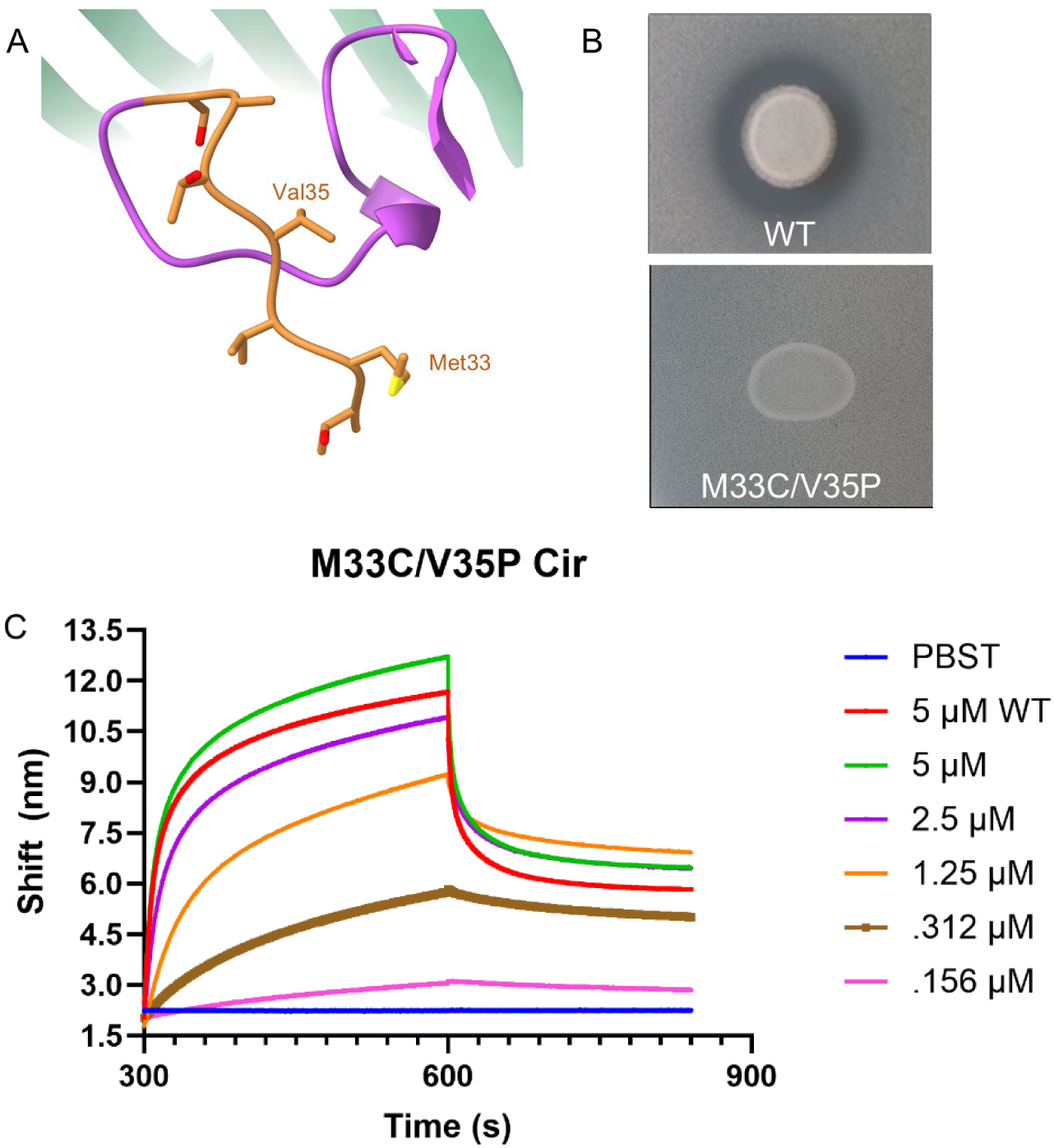
Mutation of the Cir TonB Box Inhibits MccV Killing, Not Binding. A) A zoom-in of the TonB Box within the N-Terminal domain of Cir, highlighted in orange and with the mutated residues labeled. B) Zone of inhibition assays using WT MccV secreted onto a lawn of WT or M33C/V35P Cir-expressing cells. We observed a zone of inhibition only when WT Cir was produced. C) BLI assay using WT and M33C/V35P Cir on WT MccV. 100 nM MccV was loaded to BLI sensors and they were subsequently probed with dilutions of M33C/V35P Cir, with 5 μM WT Cir included as a positive control and buffer only as a negative. Trace is representative of n=4 experiments from two independent protein preparations.

Taken together, our findings in the current study demonstrate the specific interaction between Cir and MccV, built upon a high-affinity binding event reminiscent of that observed in another Cir ligand, Colicin Ia. The flexible extracellular loops of Cir facilitate a conformational change to reveal a normally occluded region of electropositivity, and secreted MccV kills Cir-expressing *E. coli* only when this cavity is intact and when the target cell has a functioning Ton system.

## DISCUSSION

Microcins were first described as eco-active, growth-inhibitory molecules of low molecular weight produced by Gram-negatives; they were shown to resist proteases and extremes of temperature and pH^43,44^. It was during these early characterizations that the possibility of using microcins as targeted antimicrobials was considered, and while still a promising candidate, they are as-yet undeveloped and understudied^45,46^. Few microcins have been characterized to the point of having their functions confirmed, and most available information is restricted to the genetic or bioinformatic level, with the exception of Microcin D93 for which some structural data are available^47^. For MccV, the genes for microcin production and export are known and have served as the blueprint for identifying other Class II species^28,29,48^.

TBDTs have long been a focal point in understanding active transport across Gram-negative membranes, with the first examples of their crystal structures being solved in the nineteen nineties^49,50^. They play critical roles in bacterial metabolic homeostasis by facilitating the uptake of nutrients including metal ions, vitamins, and carbohydrates^9,39^. Cir is an iron regulated TBDT found in *E. coli* which coordinates iron uptake by binding catecholate siderophores such as dihydroxybenzoyl serine^51,52^, but can also be co-opted by bacteriocins during interspecies competition. The high-resolution structure of MccV bound to Cir allows us to begin detangling the molecular nature of MccV binding to target cells, and provides a critical step in evaluating microcin potential in structure-based drug design.

Targeting TBDTs for drug and vaccine development is a topic which has gained momentum over the past two decades. Recently, the β-lactam cefiderocol, which uses siderophore-like properties to access the periplasm directly via TonB-dependent transport, has shown promising efficacy against multidrug-resistant isolates, including the ESKAPE pathogen *Acinetobacter baumannii*^53–55^. Earlier such efforts utilized siderophore conjugation to the monobactam aztreonam, examples of which reached phase I trials^56,57^. The TBDTs themselves exhibit many features which make them promising targets for either drugs or vaccines: they are surface exposed; they show wide distribution in pathogenic strains; they are critical and in some cases strictly required for establishing infections^58–61^; and for vaccines specifically, can elicit protective immune responses^62,63^. Moreover, their domains and function are generally well-conserved.

MccV binds Cir atop the plug domain in a pocket normally occluded by loops 7 and 8, which hinge open to facilitate ligand entry. This is consistent with previous reports which demonstrate substrate-dependent rearrangement of extracellular loops in other TBDTs^64,65^. For Cir, an electropositive cavity revealed by this rearrangement, flanked by residues R116, R436, and R490, is key in coordinating interaction with D85 from MccV, a binding condition that is shared with Cir ligand Colicin Ia. We submitted the Cir amino acid sequence to ConSurf^66^ (https://consurf.tau.ac.il) to assess whether these residues were conserved over *E. coli* isolates (Supplementary Figure 6). R116, which sits in the highly-conserved plug domain, is maximally conserved across queries, while R436 and the adjoining C435 show moderate conservation. R490 however shows moderate variability, which one could hypothesize is the result of evolutionary pressure from bacteriocins in the intestinal chemosphere, especially considering our data which showed it to be the residue with the most profound impact on MccV binding. This prompts interesting questions regarding the evolutionary relationships between microcins and their receptors, which future studies may aim to clarify.

Mutation of residues at the Cir/MccV binding pocket resulted in clear deficiencies in bacteriolysis and binding, but it is notable that binding was only altogether eliminated in the R490A Cir variant. Similarly, while D85A MccV produced a zone of inhibition significantly smaller than seen for WT MccV, a minor zone was still visible which was absent from D85R. Considering these observations together, one must be mindful when discussing MccV binding and allow that while the charged cavity is a central player in initial recognition, the collective network of interacting residues is likely to be much larger. Similarly, an allowance must be made for the fact that our understanding of the full mechanism of MccV import remains limited. Our data are consistent with previous reports suggesting the necessity of a functional TonB^31^, an observation which is also shared for Colicin Ia^32^, but we as yet do not have a clear picture of the events which follow MccV binding at the Cir loops or whether it interacts with other targets prior to locating SdaC. This remains an interesting area of investigation, prompting questions over the requirement, for example, for MccV unfolding to traverse the Cir pore or whether other proteins are recruited during the process. It is especially interesting that MccV and Colicin Ia share a transporter in light of their considerable size difference^67^ (Supplementary Figure 7), and considering that MccV is not predicted to have a TonB box of its own as one would find in the Group B colicins like Ia. Future efforts in this area are expected to investigate this.

A final consideration involves the use of microcins as antimicrobials and how the microcins, when deployed, will interact with the host niche inhabited by their cognate pathogen. Specifically, one wonders about competition for protein binding sites between the microcin and the “intended” substrate of its target, be it a catecholate siderophore as for Cir, or otherwise. We showed that MccV binds Cir with an affinity of approximately 1 μM, but other siderophore-receptor interactions have been shown previously to be much tighter^68^. In direct competition, then, the microcins may be naturally disadvantaged by the presence of other substrates. Further studies between known microcin/receptor pairs, including MccV and Cir, should aim to assess this obstacle and consider potential protein engineering strategies that may be necessary to maximize microcin efficacy.

## METHODS

### Bacterial Strains and Plasmids

All bacterial strains, primer sequences, and plasmid maps used in this study are available upon request. The mature coding region of *E. coli cirA* (4-Met) was cloned into a modified pET-9 vector (Novagen) containing a *pelB* signal sequence followed by a 10X histidine tag and a tobacco etch virus (TEV) protease recognition site^69^ upstream of *cirA*. Cir mutants were generated in this plasmid via Q5 site-directed mutagenesis (New England Biolabs). For MccV, the mature coding region was cloned into pET-11M (Novagen) to generate a maltose binding protein (MBP) fusion. The final construct carried MBP bearing an N-terminal 6X histidine tag, followed by a TEV cleavage site and then MccV. Mutant MccV was generated from this plasmid using Q5. All plasmids were confirmed by sequencing (Psomagen).

### Protein Expression and Purification

Cir (WT and mutants) was expressed in BL21 (DE3) *E. coli* (New England Biolabs) as follows. Chemically competent cells were transformed with appropriate plasmids and selected on Luria-Bertani (LB) medium containing 50 μg/mL kanamycin. A single colony was selected and confirmed via western blot to produce his-tagged Cir, then stocked in 25% glycerol. For large scale expression, a 5 mL starter culture from stock was grown in LB + antibiotics at 37°C overnight. This starter was then used to inoculate 2 flasks containing 1 L of Terrific Broth (TB) + antibiotics, and these were grown at 20°C with 225 RPM shaking and no inducer added, instead allowing the promoter to slowly leak. Cell pellets were collected by centrifugation (10,000 x g, 10 minutes, 4°C) and resuspended in lysis buffer (50 mM Tris-HCl pH 7.5, 200 mM NaCl, 10 mM MgCl2, 10 μg/mL DNase I (Goldbio), 100 μg/mL 4-(2-aminoethyl)benzenesulfonyl fluoride hydrochloride [AEBSF, GoldBio]) at a ratio of 100 mL buffer for every 25 g of cell pellet.

Once fully resuspended, cells were lysed via two passages through an Emulsiflex C3 (Avestin) with a homogenizing pressure of ∼17,500 PSI. The lysate was then clarified by centrifugation (20,000 x g, 10 minutes, 4°C) and the supernatant collected. Lysate was mixed with 1% Triton X-100 and gently stirred for 30 minutes at room temperature. Membrane pellets were collected by centrifugation in a Beckman Type 45 Ti rotor (234,000 x g, 45 minutes, 4°C) and supernatant was discarded. Membranes were resuspended in 1X phosphate buffered saline (PBS) using a Dounce homogenizer. This solution was mixed with an excess volume of purified MccV (described below) for 2h, then final concentrations of 40 mM imidazole and 1% DDM were added. This mixture was gently mixed for 2 hours at room temperature to solubilize the membranes. After solubilization, the solution was clarified by centrifugation in a Beckman Type 70 Ti rotor (310,000 x g, 45 minutes, 4°C) and the soluble fraction was collected. This was filtered through a 0.22 μm filter then applied to an AKTA purifier (GE) for nickel affinity chromatography. The sample was looped ten times over a 5 mL HisTrap column (Cytiva) that was equilibrated with PBS pH 7.5 plus 0.1% DDM prior to sample application. His-tagged Cir was eluted by stepwise addition of imidazole in the same buffer, with the majority eluting at 300 mM imidazole.

Fractions were analyzed by SDS-PAGE and the cleanest were carried forward. These fractions were pooled and mixed with a threefold mass excess of NAPol (Anatrace) and gently rocked at room temperature for roughly 1 h. DDM was removed by addition of freshly-hydrated Bio-Beads SM-2 (Bio-Rad) and gentle rocking for 1 h. The beads were removed by filtering through a 0.22 μm filter and the resulting protein solution was concentrated to a final volume of 500 μL using a 50 kDa cutoff centrifugal filter (Millipore). This sample was applied to a Superdex 200 Increase 10/300 GL column (Cytiva) equilibrated with 1X PBS for size exclusion chromatography. Peak fractions were again analyzed by SDS-PAGE, and sample homogeneity from the cleanest fractions was further confirmed by mass photometry^70^ using a OneMP mass photometer (Refeyn).

MccV (WT and mutant) was expressed in BL21 (DE3) *E. coli* as follows. Chemically competent cells were transformed and selected on LB + 50 μg/mL, then stocked as described above. A 5 mL starter culture was grown overnight as described and used to inoculate a flask containing 1 L 2xYT medium + antibiotics. This culture was grown at 37°C at 225 RPM shaking until the it reached an OD600 of 0.6 – 0.8, at which point protein expression was induced by addition of 1 mM Isopropyl β-D-1-thiogalactopyranoside (IPTG). Protein expression continued for 3 h before cell pellets were collected by centrifugation and resuspended in lysis buffer as described above. Cells were lysed via 5 minutes of looping through the Emulsiflex C3 and then clarified by centrifugation. The lysate was applied to an AKTA purifier and loaded onto a 5 mL HisTrap column in 1X PBS to capture the His-tagged MBP fusion partner. MBP_MccV eluted at a range from 50 to 300 mM imidazole in 1X PBS. Fractions were subjected to SDS-PAGE and relevant fractions were pooled and mixed with 2 mM dithiothreitol (DTT) and 2 mg of His-tagged TEV protease which had been purified as described elsewhere^71^. The protein/TEV mixture was pipetted into pre-hydrated 3 kDa cutoff dialysis tubing (Millipore) and dialyzed against an excess of 1X PBS for 48 h at 4°C while TEV cleavage of the MBP partner proceeded. After dialysis, the resulting solution was passed again over a nickel column to recapture the his-tagged TEV and MBP, and the flowthrough was collected. The remaining purified, untagged MccV was concentrated in a 3 kDa cutoff centrifugal filter and its purity assessed by SDS-PAGE.

### Cryo-EM Sample Preparation and Data Acquisition

3 μL of Cir/MccV complex (7.8 mg/mL) was applied to a Quantifoil R 1.2/1.3 300 mesh cryo-EM grid (Protochips, Inc.) that had been glow discharged for 45 seconds at 15 mA prior to sample application. The grid was blotted and plunged immediately into liquid ethane using a Vitrobot Mark IV (ThermoFisher). Blotting conditions were as follows: 4°C, 100% humidity, blot force +5, blot time 3-7 seconds, wait time 0 seconds. EM data were collected using SerialEM^72^ on a Titan Krios G3 (ThermoFisher) equipped with an Imaging Filter Quantum LS and a K3 direct electron detector (Gatan) with an acceleration voltage of 300 keV. The calibrated pixel size was 0.415 Å/pixel in super-resolution mode. A total of 7,606 dose-fractionated movies were collected with acquisition software operating in counting mode. The total electron dose was 60 e/Å^2^ over 30 frames, and a nominal defocus range of -0.8 to -2.4 μm.

### Data Processing and Model Building

All data processing was performed in CryoSPARC v4^73^. Movies were imported then gain and motion corrected using patch motion correction with a binning factor of two (0.83 Å/pixel). Patch CTF was used to estimate CTF parameters. Motion corrected frames were curated and 5,960 exposures were retained based on CTF fit resolution and relative ice thickness. A blob picker was used on 500 exposures and identified ∼1.1 million particles which were extracted (bin 2) and submitted to 2D classification. 134,798 particles were selected and used to generate two Ab-Initio classes, while the rejected particles were used to generate four “junk” Ab-Initio classes. The 2-class Ab-Initio job was plugged immediately into heterogeneous refinement and the better class was selected based on evidence of TBDT domains (β-barrel, plug, loops, micelle). This 3D volume was used to generate templates, which were used in a template picker job on the entire 5,960 micrograph dataset, resulting in ∼7.7m extracted particles (bin 2). These particles were plugged into a heterogeneous refinement using the 3D volume which was used for template generation alongside the four “junk” classes. The best class was carried forward and used in Ab-Initio generation, with the top class again selected for heterogeneous refinement alongside the “junk” volumes. This process was repeated three times to remove as many junk particles as possible, then a final 2D classification was performed on the best remaining class to leave 304,879 particles, which were then re-extracted at box size 256 x 256. These particles and volume were added to a non-uniform refinement, and the resulting volume had its hand flipped by homogenous reconstruction. The particles were submitted to global CTF refinement, and then another non-uniform refinement using the hand-flipped volume. The particles from this volume were then rebalanced to ensure equal representation of viewing directions, and the remaining 102,419 particles were submitted to local refinement using a tight-fitting mask to omit the micelle, yielding a 2.9 Å final density map. A schematic representation of this workflow is shown in Supplementary Figure 2.

An initial predicted model for Cir/MccV was generated in AlphaFold2^38^ using the multimer function, which was then docked into the EM density map using PHENIX^74^. The placed model was then manually adjusted in Coot^75^, using a DeepEMhancer^76^ map to aid visualization of density fit and rotamer assignments; local real-space refinement and regularization were performed against the experimental map. Final refinement and validation of the model was performed in PHENIX with additional relaxing in ISOLDE^77^.

### Biolayer Interferometry

BLI experiments were performed using a Gator Plus automated BLI instrument (Gator Bio). Purified MccV (WT and mutants) was biotinylated using an EZ-Link Sulfo-NHS-LC biotinylation kit (ThermoFisher) according to the manufacturer’s protocol. Prior to experiments, streptavidin-coated SA-XT sensors (Gator Bio) were soaked in 300 μL PST-T in a 96-well MAX plate. In the instrument, sensors were equilibrated to baseline in PBS-T for 60 seconds, then MccV-biotin was loaded at a concentration of 100 nM. Loading time was 120 seconds. Loaded sensors were allowed to return to baseline for 120 seconds before the association phase. For association, Cir (WT and mutants) was serial twofold diluted in PBS-T (5 μM -> 78 nM), and these dilutions were added to the sensors for an association time of 300 seconds. After association, sensors were returned to PBS-T for 240 seconds of dissociation time. For experiments containing mutated Cir or MccV, a probe loaded with WT MccV and addition of 5 μM WT Cir was added as a positive control. Addition of PBS-T alone was used as a negative control. Kinetic analysis was performed in the Gator Navigator Software, and raw response data were plotted in GraphPad Prism for visualization.

### Sedimentation Velocity Analytical Ultracentrifugation

NAPol solubilized Cir and its mixtures with MccV were analyzed by SV-AUC in phosphate-buffered saline. Sedimentation velocity experiments were conducted at 50,000 rpm (198,800 x g at 7.1 cm) and 20°C on a Beckman Coulter ProteomeLab XL-I analytical ultracentrifuge and An-50 Ti rotor. Sedimentation data were time-corrected and analyzed in SEDFIT^78^ in terms of a continuous c(*s*) distribution of Lamm equation solutions. Solution densities ρ, solution viscosities η, and protein partial specific volumes were calculated in SEDNTERP^79^. The protein refractive index increment was calculated in SEDFIT. The partial specific volume for NAPol was calculated based on its chemical composition following the method of Durchschlag and Zipper^80^, and a refractive index increment of 0.149 cm^3^g^-1^ was used. Absorbance and interference c(*s*) distributions were analyzed simultaneously using the fitted f/fo membrane protein calculation module in GUSSI^81^ to obtain the protein and amphipol contributions to the sedimenting complex of interest.

### Flow Cytometry

MccV RBD with an N-terminal dansyl modification was synthesized by GenScript. Strains were grown in LB media with appropriate antibiotics at 37°C to exponential phase (0.2% arabinose was added to cultures containing pBAD plasmids encoding *cirA*). Cells were pelleted (4,000 RPM, 10 minutes), supernatants were removed, and cells were gently resuspended in 1X PBS containing 50 mM glucose to a final cell density of OD600=0.2. For each condition tested, 1 mL of cell suspension was added to a 5 mL polystyrene round-bottom tube (Corning). For cells treated with MccV RBD-dansyl, resuspended peptides were added to a final concentration of 53 µM. Tubes were incubated in the dark for 30 minutes, then analyzed using a Sony SA3800 spectral cell analyzer until 10,000 events were captured. Results were analyzed, gated, and visualized using FlowJo software (v 10.10.0).

### MccV Bactericidal Assays

The WT *cirA* sequence with an appended C-terminal V5 tag was cloned into a pBAD vector with ampicillin resistance using standard cloning techniques. WT MccV and its associated immunity protein sequence were previously cloned into a pBAD plasmid with kanamycin resistance. All Cir and MccV residue mutants were constructed using QuikChange Site-Directed Mutagenesis kit (Agilent). Cir mutant plasmids were used to transform a *cirA*::kan strain from the Keio collection in order to isolate the sensitivity of the Cir variant to MccV. Plasmids encoding MccV were used to transform an *E. coli* strain containing the microcin secretion system on plasmid pACYC.

Bactericidal activity was assayed using zone of inhibition assays. All strains were grown overnight at 37°C in LB media with appropriate antibiotics. Prey strain cultures were pelleted (4,000 RPM, 10 minutes), supernatants were removed, and cells were resuspended in LB media. Each prey strain was added to 10 mL of LB containing 0.75% agar and 0.2% arabinose to a final OD600 of 0.001. The mixture was poured over a solidified bottom layer of LB containing 1.5% agar. Overnight cultures of bacteria containing the microcin secretion system and MccV plasmids were normalized to the same OD600 in 500 µL, then pelleted (4,000 RPM, 10 minutes). Cell pellets were resuspended in 50 µL of LB and 5 µL of predator strain was spotted onto plates containing the prey strains. Plates were grown overnight at 37°C and zones of inhibition were inspected to infer mutant sensitivity.

### Expression Test for Cir Mutants

To ensure that variability in prey mutant sensitivity was not due to differences in protein expression, Cir production was assayed by western blot. Strains were grown overnight at 37°C in LB with appropriate antibiotics and 0.2% arabinose. The following day, 1.5 mL of overnight culture was pelleted and resuspended in 1 mL of 10 mM Tris pH 7.3 and 150 mM NaCl. All cultures were then normalized to the same density and 20 µL of cell suspension was mixed with SDS loading buffer and incubated at 99°C for 10 minutes. Samples were loaded into a NuPAGE Bis-Tris 4-12% gel (Invitrogen) and run in MES buffer. Proteins were then transferred onto a PVDF transfer membrane with 0.45 µm pore size (Thermo Scientific), according to the manufacturer’s instructions. After transfer, membranes were incubated in blocking buffer (1X TBS, 0.1% Tween, 5% milk) for one hour at room temperature, then incubated overnight at 4°C in blocking buffer containing anti-V5 tag antibody (1:5000 ratio). The membrane was washed three times in blocking buffer (10 minutes each) and incubated with blocking buffer containing anti-rabbit-HRP antibodies (1:1500 ratio) for four hours at 4°C. After incubation, the membrane was washed three times in TBST (1X TBS, 0.1% Tween). Membranes were briefly incubated with SuperSignal West Femto Maximum Sensitivity Substrate (Thermo Scientific) and imaged on a BioRad ChemiDoc imaging system.

## ACKNOWLEDGEMENTS

This study utilized resources from the NIH Multi-Institute Cryo-EM Facility (MICEF), the National Heart, Lung, and Blood Institute (NHLBI) Biophysics Core, and the NIH HPC Biowulf cluster (http://hpc.nih.gov). The authors would like to thank Yanxiang Cui, Huaibin Wang, Ulrich Baxa, and Bertram Canagarajah for microscopy technical support and data management, and Grzegorz Piszczek and Di Wu for biophysics core support. SM, IB, RG, and SB are supported by the Intramural Research Program of the National Institute of Diabetes and Digestive and Kidney Diseases. AO and BD are supported by the National Institutes of Health (R01 AI182365, R56 AI179199), Welch Foundation F-2137, and Army Research Office W911NF2010195.

