## Supplementary Information for "Structural Insights into Cir-mediated Killing by the Antimicrobial Protein Microcin V"

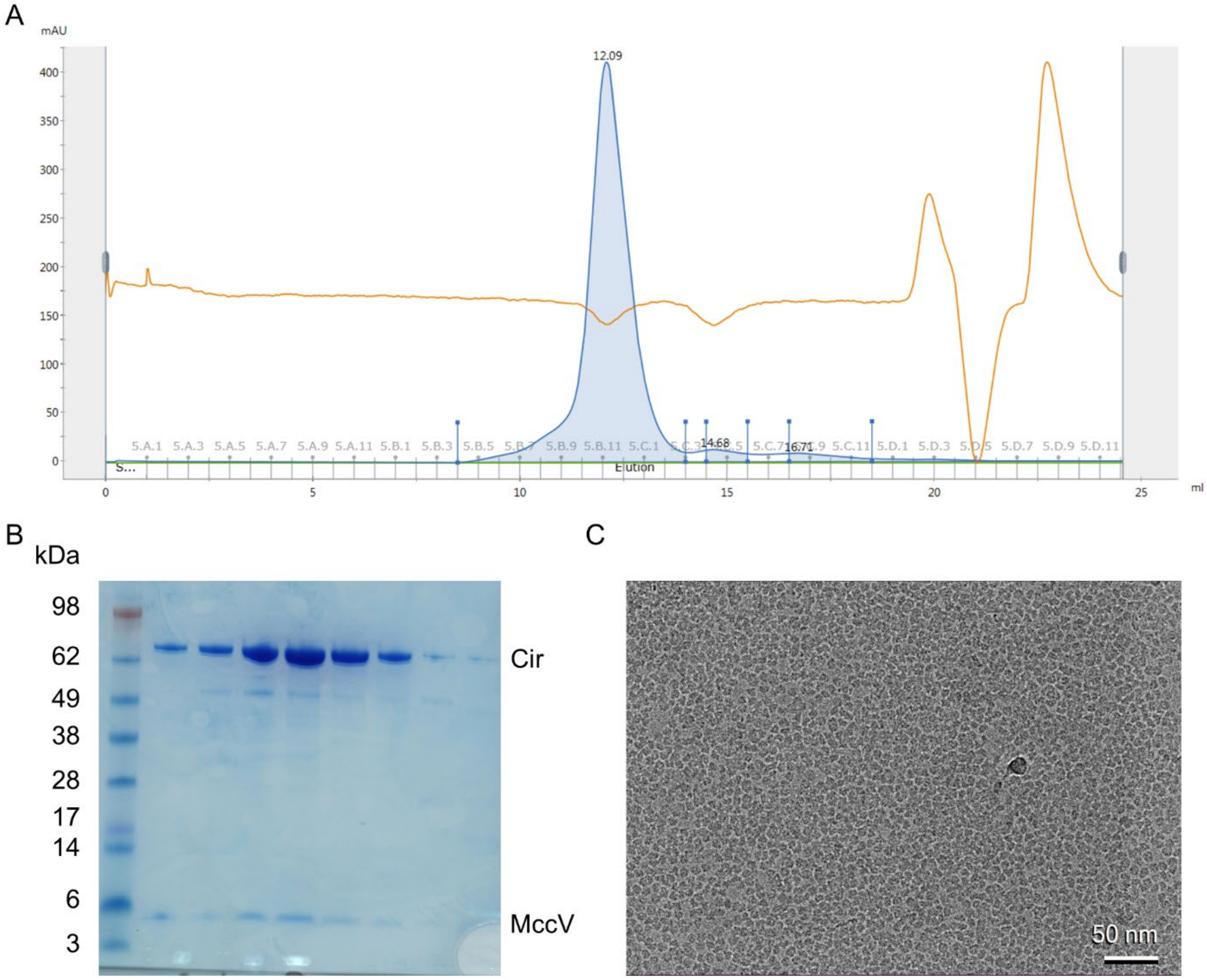


**Supplementary Figure 1 – Sample Purity of Cir/MccV Complex.** A) Size exclusion chromatography elution profile for Cir/MccV complex showing protein eluting as a clean, single peak. B) Coomassie stained protein gel showing relevant fractions from panel A. Fractions contained both Cir and MccV. C) Representative micrograph from cryo-EM data collection showing particle density and distribution in thin ice. Sample was loaded at 7.8 mg/mL.

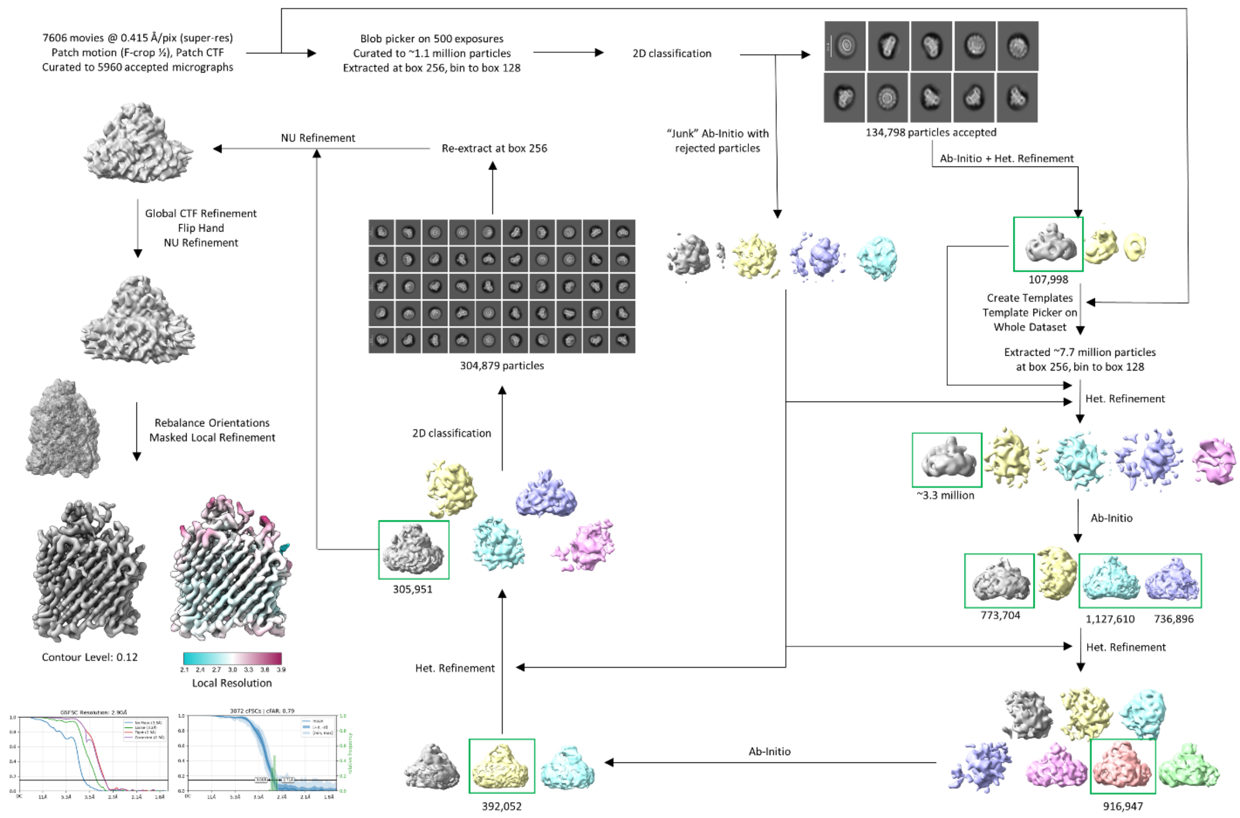


**Supplementary Figure 2 –** Schematic diagram of cryo-EM image reconstruction workflow for Cir/MccV RBD complex, resulting in a 2.9 Å consensus map.

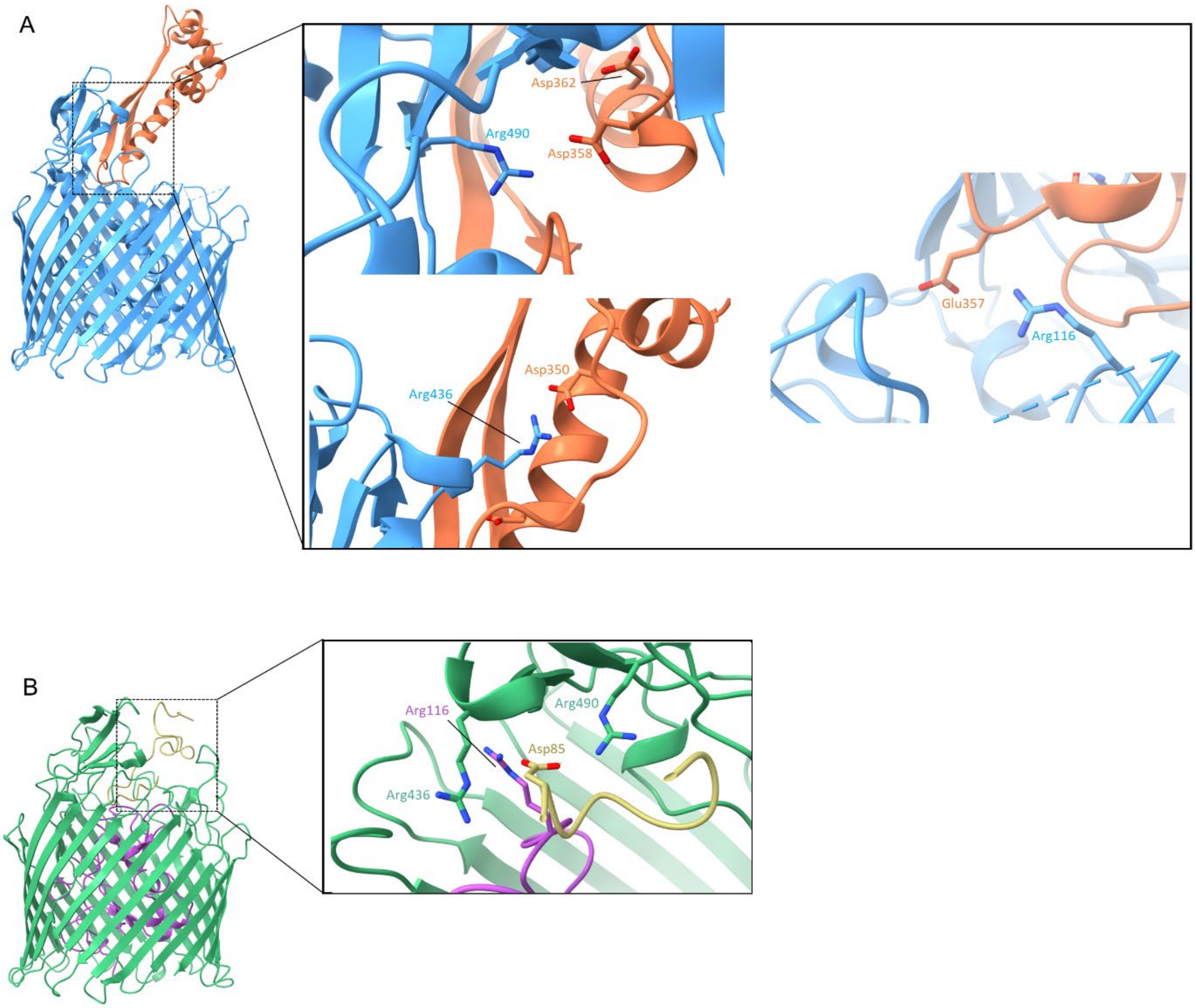


**Supplementary Figure 3 – Comparison of Cir Interacting Residues with Colicin Ia and MccV.** A) Structure of Cir liganded by the receptor binding domain of Colicin Ia (PDB code 2HDI). Zoomed box shows the three positively charged arginine residues corresponding to those described for the Cir/MccV structure in figure 4. B) Cir/MccV structure and zoom of charged residues for comparison.

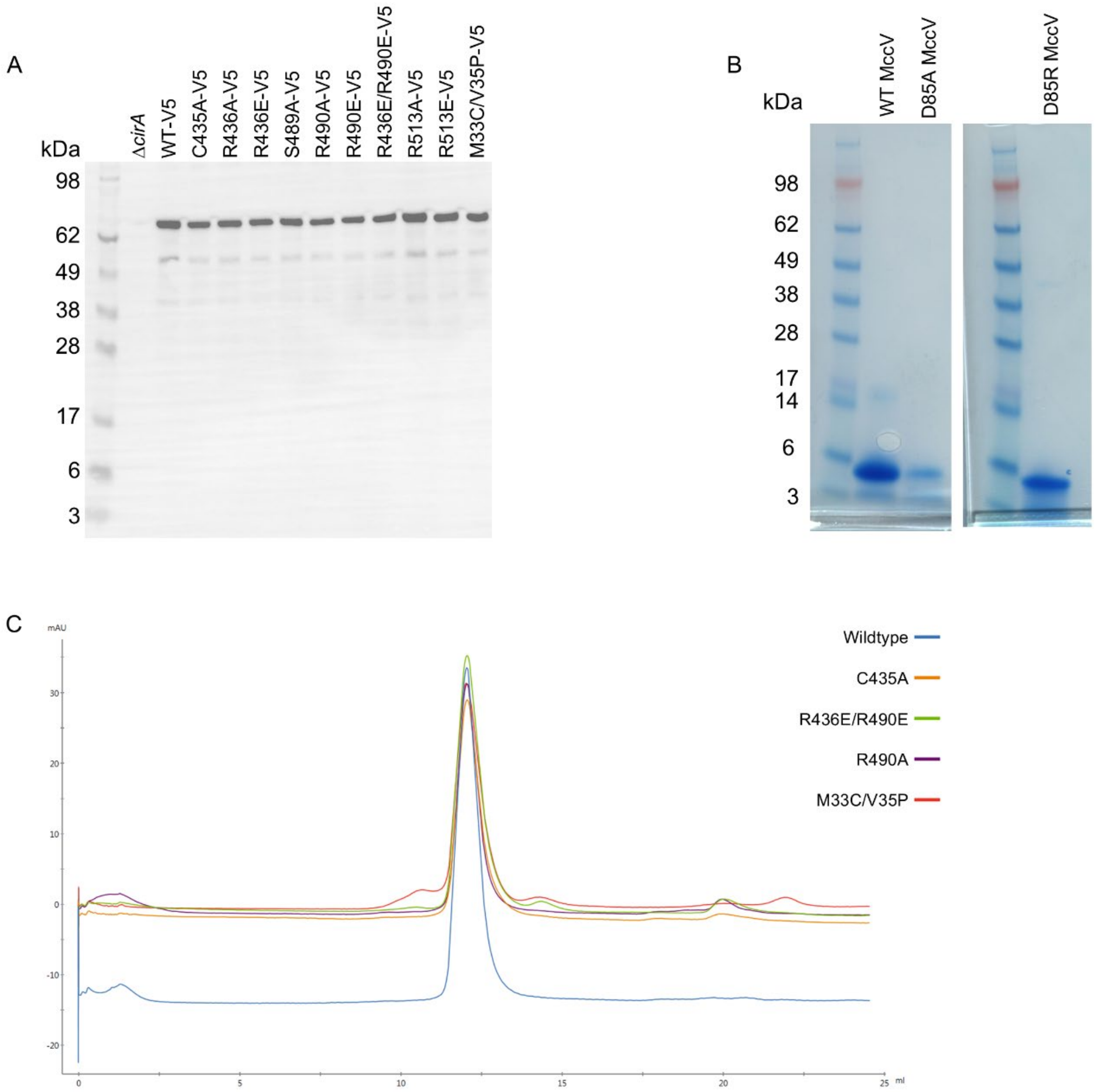


**Supplementary Figure 4 – Cir Mutants Are Stably Expressed and Purified.** A) Western blot showing comparable levels of Cir mutant expression and lack of detectable degradation. Proteins were detected using C-terminal V5 tags. B) Coomassie stained protein gels of WT, D85A, and D85R MccV. C) Overlay of size exclusion chromatography elution traces for WT, C435A, R490A, R436E/R490E, and M33C/V35P Cir. Proteins all eluted at the same retention volume and as clean, single peaks.

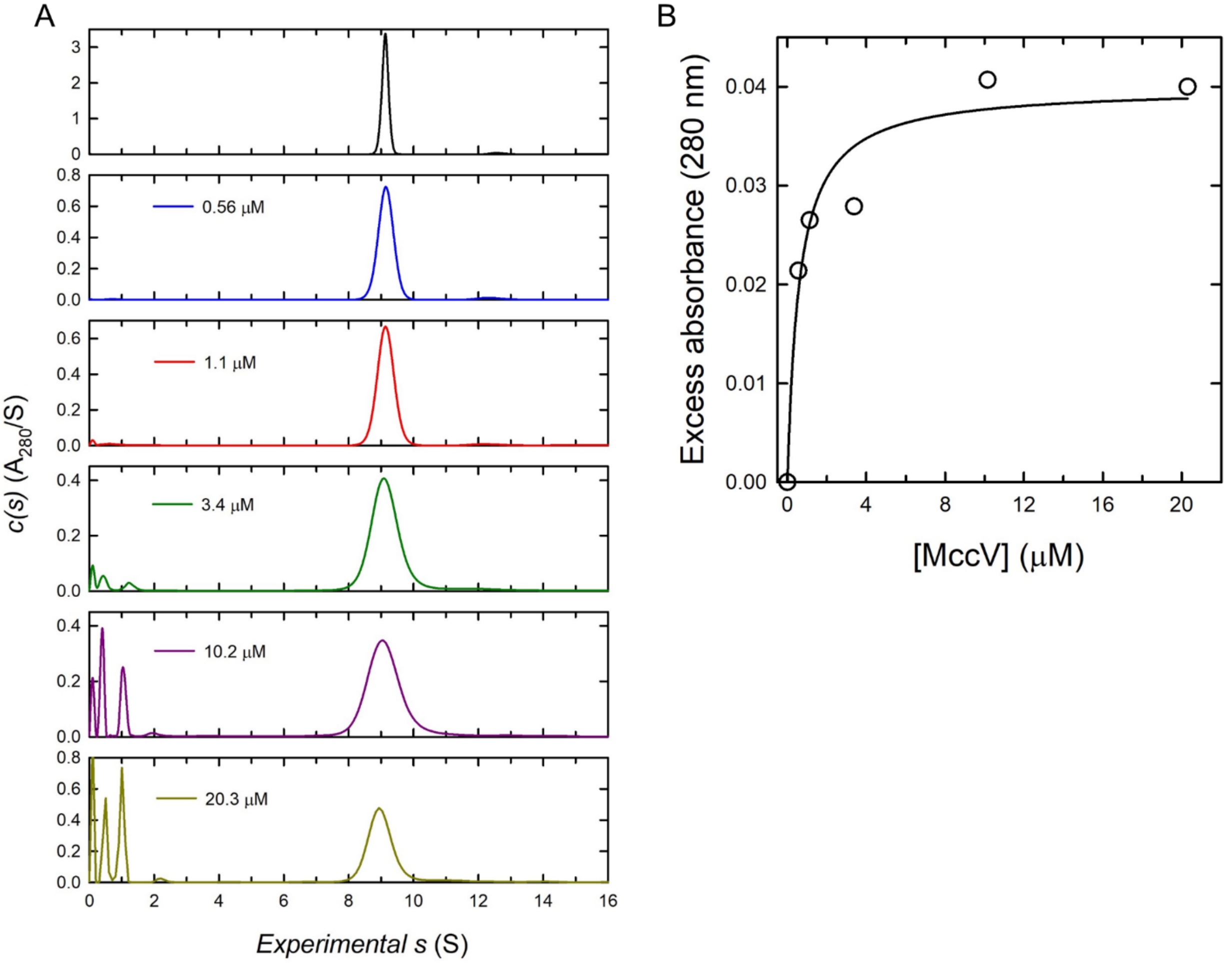


**Supplementary Figure 5 – Analytical Ultracentrifugation Analysis of MccV Binding to Cir.** A) Absorbance sedimentation c(*s*) profiles for Cir. Top panel - Absorbance sedimentation c(*s*) profile for NAPol solubilized Cir at 5.6 μM showing a species at 9.11 S. A membrane protein analysis returns a protein mass of 75 ± 13 kDa, supporting a Cir monomer, and a complex mass of 195 ± 41 kDa. The excess mass represents the NAPol contribution. Lower Panels - Absorbance sedimentation c(*s*) profiles for NAPol solubilized Cir at 2.7 μM in the presence of MccV. The concentration of MccV added is indicated on each plot. The absorbance signal for the Cir species provides a measure of MccV binding, which was obtained by integration. B) Binding of MccV to Cir. Excess absorbance of the Cir c(*s*) absorbance contribution, indicating MccV binding, is plotted as a function of the MccV concentration. Data were fit to a single binding model to obtain a dissociation constant KD of 0.6 ± 0.2 μM. The maximum excess absorbance at high MccV concentrations corresponds to the value expected when a single MccV binds to 2.7 μM of Cir, validating the 1:1 binding.

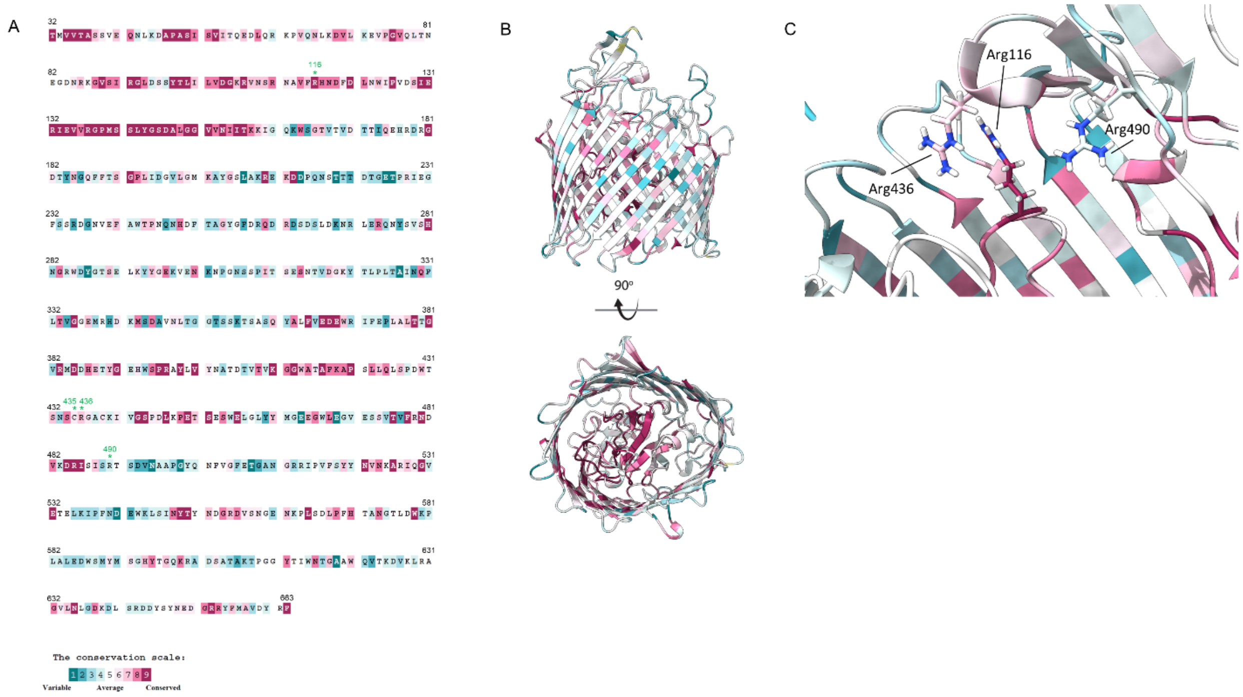


**Supplementary Figure 6 – Amino Acid Sequence Conservation of Cir.** A) The Cir amino acid sequence (beginning at mature start) was submitted to the ConSURF server for prediction of conserved functional regions, with a focus on Arg 116, Cys 435, Arg 436, and Arg 490. The sequence is colored according to the conservation scale shown at the bottom, and the relevant residues are highlighted with a green asterisk. B) Cir/MccV model colored according to conservation data in A. C) Conservation of R116, R436, and R490.

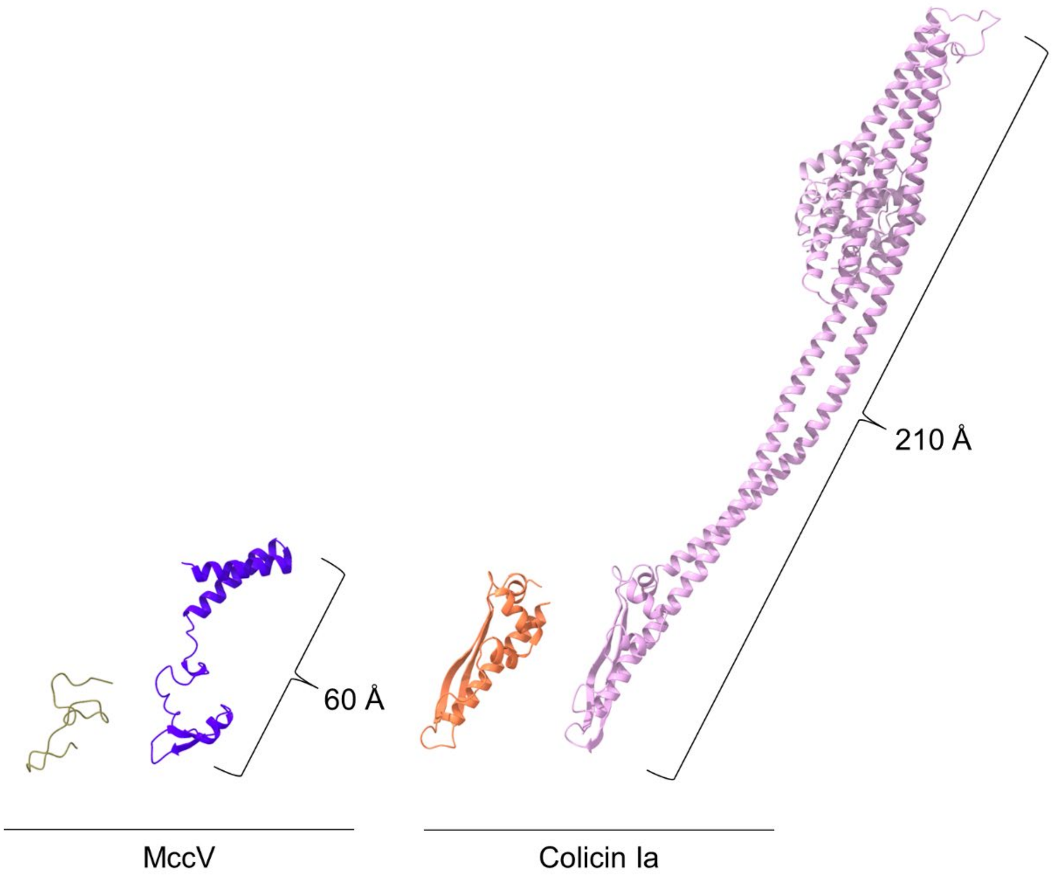


**Supplementary Figure 7 – Size Comparison of MccV and Colicin Ia.** Structures of (left to right) the MccV RBD, mature MccV (AlphaFold2 prediction), Colicin Ia RBD (from PDB 2HDI), and whole Colicin Ia (PDB 1CII).
