## Supplementary material for "Structural Insights into Cir-mediated Killing by the Antimicrobial Protein Microcin V": Table 1

### Table 1: Cryo-EM Data Collection and Structure Statistics

| | Cir/MccV |
|---|---|
| ***Data Collection*** | |
| Magnification | 105,000x |
| Voltage (keV) | 300 |
| Total Dose (e/Å$^2$) | 59.65 |
| Number of Frames | 30 |
| Exposure Time (s/frame) | 0.075 |
| Defocus Range (μm) | -0.8 to -2.4 |
| Pixel Size (Å) | 0.83 |
| ***Image Processing*** | |
| Movies Collected | 7,606 |
| Micrographs Selected | 5,960 |
| Particles Extracted | 7,726,727 |
| Final Map Particles | 102,419 |
| Symmetry Imposed | C1 |
| FSC Threshold | 0.143 |
| Final Map Resolution (Å) | 2.9 |
| Resolution Range (Å) | 3.06 – 2.71 |
| ***Atomic Model*** | |
| Protein Residues | 664 |
| Chains | 2 |
| ***Validation*** | |
| Ramachandran Favored (%) | 97.73 |
| Ramachandran Allowed (%) | 2.27 |
| Ramachandran Outliers (%) | 0 |
| Rotamer Outliers (%) | 0 |
| RMSD Bond Lengths (Å) (# > 4σ) | 0.002 (0) |
| RMSD Bond Angles (°) (# > 4σ) | 0.419 (0) |
| Clash Score | 5.60 |
| Map CC (mask) | 0.87 |
| Map CC (volume) | 0.88 |
| ***Deposition IDs*** | |
| PBD | 9NN6 |
| EMDB | EMD-49565 |
